## Supplementary Information for "Global diversity of soil-transmitted helminths reveals population-biased genetic variation that impacts diagnostic targets"

Origin of samples and data	2

Supplementary Fig. 1: Comparison of helminth-positive samples with single or multiple helminth species infections.	7

Supplementary Fig. 2: Geographic distribution of helminth-positive samples from faecal and worm isolates.	9

Supplementary Fig. 3: Exploratory analysis of population and genetic structure of samples positive for *Ascaris lumbricoides*.	10

Supplementary Figure 4. Preferential mapping of samples positive for *Ascaris* spp. to one of three *Ascaris* reference genomes from geographically distinct isolates.	12

Supplementary Figure 5. Resolved genetic analyses of *Ascaris* spp. positive samples mapping to *Ascaris suum*.	14

Supplementary Figure 6. Population structure and genetic differentiation between *Trichuris* *trichiura* populations.	16

Supplementary Figure 7. Distribution of nuclear repeat and ribosomal sequences used as diagnostic targets in the genome assemblies.	19

Supplementary Figure 8. Comparison of relative copy-number of nuclear repeats and mitochondrial genomes based on sequencing coverage.	20

Supplementary Figure 9. Assessing the presence, distribution, and impact of genetic variation within diagnostic qPCR targets of *Trichuris trichiura*.	22

References	24

### Origin of samples and data

We aimed to source worm isolates and egg/faecal data with strong infections for any soil-transmitted helminth (STH) present. Below is a breakdown of the number and types of samples per country, any relevant ethical approvals, and accession numbers relating to publicly available data.

a. Argentina (n = 1).
   i. n = 1; whole genome sequencing data of DNA extracted from concentrated pools of eggs of all STH species; the original study protocol was approved by the bioethics committee of Colegio de Médicos de la Provincia de Salta and the IRBs of BCM (protocol number H-34926); citation: this study. Key collaborators: Nicolas R Caro, Ruben O Cimino, Alejandro J Krolewiecki, Rojelio Mejia.
b. Bangladesh (n = 10).
   i. n = 10; whole genome sequencing data of DNA extracted from faecal samples; the original study protocol was approved by the Ethical Review Committee at icddr,b (PR-14105), the Committee for the Protection of Human Subjects at the University of California, Berkeley (2014-08-6658), and the institutional review board at Stanford University (27864); citation: this study. Key collaborators: John M Colford Jr, Jade Benjamin-Chung, Steven A Williams.
c. Benin (n = 25).
   i. n = 25; whole genome sequencing data of DNA extracted from faecal samples, from the DeWorm3 project. The original study protocol was approved by the Institut de Recherche Clinique au Bénin (IRCB) through the National Ethics Committee for Health Research (002-2017/CNERS-MS) from the Ministry of Health, the Human Subjects Division at the University of Washington (STUDY00000180) and the Data Safety and Monitoring Committee (DSMC); citation: this study. Key collaborators: Moudachirou Ibikounlé, Adrian JF Luty, Judd L Walson.
d. Cameroon (n = 6).
   i. n = 1; whole genome sequencing data of DNA extracted from a *S. mansoni* worm; ENA study accession: [PRJEB2679](https://www.ebi.ac.uk/ena/browser/view/PRJEB2679); Sample accession code: ERR103050; citation: ^1^.
   ii. n = 5, whole genome sequencing data of DNA extracted from concentrated eggs from faecal samples; ENA study accession: PRJEB44010; sample accession codes: ERR9805789-93; citation: ^2^.
e. China (n = 8)
   i. n = 1, whole genome sequencing data of DNA extracted from an *N. americanus* worm*;* ENA study accession: [PRJNA304165](https://www.ebi.ac.uk/ena/browser/view/PRJNA304165); sample accession code: SRR2968128; citation: ^3^.
   ii. n = 7, whole genome sequencing data of DNA extracted from *T. trichiura* worms; ENA study accession: PRJEB44010; sample accession codes: ERR9805779-785; citation: ^2^.
f. Democratic Republic of Congo (n = 2).
   i. n = 1, whole genome sequencing data of DNA extracted from samples ; citation: ^4^.
   ii. n = 1, whole genome sequencing data of DNA extracted from a faecal extract, part of the same protocol approval above. The original study protocol was approved by the Democratic Republic of Congo / University Hospital, Ghent University, Belgium (M104; Catholic University of Bukavu, Democratic Republic of the Congo [Ref: UCB/CIE/NC/016/2016], the Ministry of Public Health, Democratic Republic of the Congo [Ref: 062/CD/DPS/SK/2017]); citation: this study.
g. Ecuador (n = 1).
   i. n = 1, whole genome sequencing data of DNA extracted from an *A. lumbricoides* worm, ENA study accession: [PRJNA304165](https://www.ebi.ac.uk/ena/browser/view/PRJNA304165); Sample accession code: SRR2968217; citation: ^3^.
h. Ethiopia (n = 5).
   i. n = 2, whole genome sequencing data of DNA extracted from faecal samples; ETH_ET018, ETH_ET103 in Papaiakovou et al (2023); ENA study accession: [PRJNA847183](https://www.ebi.ac.uk/ena/browser/view/PRJNA847183); sample accession codes: SAMN28922051-SAMN28922052; citation: ^4^.
   ii. n = 2, whole genome sequencing data of DNA extracted from two *A. lumbricoides* worms. The original study protocol was approved by Ethical Review Committee, Faculty of Medicine and Health Sciences / University Hospital, Ghent University, Belgium [Ref: B670201627755 and PA2014/003], Jimma University, Ethiopia (Ref: RPGC/547/2016); citation: this study. Key collaborators: Piet Cools, Bruno Levecke, Zeleke Mekonnen.
   iii. n = 1; whole genome sequencing data of DNA extracted from a faecal sample. The original study protocol was approved by the Ethical Review Committee, Faculty of Medicine and Health Sciences / University Hospital, Ghent University, Belgium (Ref: B670201627755 and PA2014/003), and by the Jimma University, Ethiopia (Ref: RPGC/547/2016); citation: this study. Key collaborators: Piet Cools, Bruno Levecke, Zeleke Mekonnen.
i. Fiji (n = 2).
   i. n = 2, whole genome sequencing data of two aliquots of DNA extracted from a single *A. lumbricoides* worm; the worm was provided by the Natural History Museum, London, UK under registration number: 2012.11.19.1, *Ascaris lumbricoides* Linnaeus, 1758 -- Ascaridinae; Ascarididae; Ascaridoidea; Spirurina; Rhabditida; Chromadorea, 1, spirit material.
j. Guadeloupe (n = 4).
   i. n = 4, whole genome sequencing data of DNA extracted from *S. mansoni* worms; ENA study accession: [PRJEB3054](https://www.ebi.ac.uk/ena/browser/view/PRJEB3054); Sample accession codes: ERR539842-45; citation: ^1^.
k. Honduras (n = 8).
   i. n = 8, whole genome sequencing data of DNA extracted from *T. trichiura* worms; ENA study accession: PRJEB44010; sample accession codes: ERR9805798-805; citation: ^2^.
l. India (n = 25).
   i. n = 25, whole genome sequencing data of DNA extracted from faecal samples, part of the DeWorm3 project. The original study protocol was approved by the Christian Medical College Institutional Review Board in Vellore, India (10392). The study was also approved by The Human Subjects Division at the University of Washington (STUDY00000180) and the genome skimming project was revised by the Data Safety and Monitoring Committee (DSMC); citation: this study. Key collaborators: Sitara SR Ajjampur, Malathi Manuel, Judd L Walson.
m. Italy (n = 2).
   i. n = 2; whole genome sequencing data of DNA extracted from faecal samples; CAM1 & CAM2 in Papaiakovou et al (2023); ENA study accession: PRJNA847183; sample accession code: SAMN28922036-37; citation: ^4,5^.
n. Kenya (n = 76)
   i. n = 68, whole genome sequencing data of DNA extracted from individual *A. lumbricoides* worms; ENA study accession: PRJNA511012; sample accession codes: SRX5228374-SRX5228441; citation: ^6^.
   ii. n = 7; whole genome sequencing data of DNA extracted from faecal samples; the original study protocol was approved by the Scientific and Ethics Review Committees (ERC) of the Kenya Medical Research Institute (KEMRI, SSC #1820); citation: (Onkanga *et al.*, 2016; Secor *et al.*, 2020). Key collaborators: Maurice Odiere, Pauline Mwinzi.
   iii. n = 1; whole genome sequencing data of DNA extracted from a *S. mansoni* worm; ENA study accession: PRJEB2679; citation: ^1^.
o. Malawi (n = 25).
   i. n = 25; whole genome sequencing data of DNA extracted from faecal samples, part of the DeWorm3 project. The original study protocol was approved by The London School of Hygiene and Tropical Medicine (12013), The College of Medicine Research Ethics Committee (P.04/17/2161) in Malawi. The study was also approved by The Human Subjects Division at the University of Washington (STUDY00000180) and the genome skimming project was revised by the Data Safety and Monitoring Committee (DSMC); citation: this study. Key collaborators: Robin Bailey, David Chaima, Khumbo Kalua, Judd L Walson, Stefan Witek-McManus.
p. Malaysia (n = 650).
   i. n = 650; whole genome sequencing data of DNA extracted from faecal samples; ENA study accession: [PRJNA797994](https://www.ebi.ac.uk/ena/browser/view/PRJNA797994); sample accession codes: SAMN25042866-25043515; citation: ^7^.
q. Mozambique (n = 20).
   i. n = 20; whole genome sequencing data of DNA extracted from concentrated eggs from faecal samples; the original study protocol (WASH-IT) was approved by the National Bioethics Committee for Health in Mozambique. ENA study accession: PRJEB53235; citation: this study. Key collaborators: Maria Cambra-Pellejà, Anélsio Cossa, Javier Gandasegui, Berta Grau-Pujol, Inácio Mandomando, Maria Martínez-Valladares, Augusto Messa Jr., Osvaldo Muchisse, Jose Muñoz, Valdemiro Novela, Charfudin Sacoor.
r. Myanmar (n = 38)
   i. n = 6; whole genome sequencing data of DNA extracted from faecal samples: SAMN28922044 (MMR_TKU23), SAMN28922045 (MMR_TKU25), SAMN28922049 (MMR_TKU102), SAMN28922046 (MMR_NDK63), SAMN28922047 (MMR_NDK92), SAMN28922048 (MMR_NDK113) in Papaiakovou et al (2023); Study accession number: [PRJNA847183](https://www.ncbi.nlm.nih.gov/bioproject/847183); citation: ^4^.
   ii. n = 32; whole genome sequencing data of DNA extracted from faecal samples; the original study protocol was approved by Imperial College London, UK (Ethical Review Ref: 17IC4249 and 17IC4249 NoA1); citation: this study. Key collaborators: Roy M Anderson, Julia Dunn.
s. Nigeria (n = 11).
   i. n = 11; whole genome sequencing data of DNA extracted from faecal samples; the original study protocol was approved by the Health Research Ethics Committee of the Kebbi State Ministry of Health, Nigeria (reference number:105:23/2021); citation: this study. Key collaborator: Olumide Ajibola.
t. Puerto Rico (n = 1).
   i. n = 1; whole genome sequencing data from a *S. mansoni* worm; ENA study accession: PRJEB2679; sample accession code: ERR046038; citation: ^1^.
u. Senegal (n = 1).
   i. n = 1; whole genome sequencing data of DNA extracted from an *S. mansoni* worm; ENA study accession: PRJEB2679; sample accession code: ERR103049; citation: ^1^.
v. South Africa (n = 7)
   i. n = 7; whole genome sequencing data of DNA extracted from faecal samples; the original study protocol was approved by the Biomedical Research Ethics Administration, University of KwaZulu-Natal KZN (Ref BF029/07); citation: this study. Key collaborator: Eyrun F Kjetland.
w. Sri Lanka (n = 7).
   i. n = 6; whole genome sequencing data of DNA extracted from faecal samples; ENA study accession: PRJNA847183; sample accession codes: SAMN28922038-43; citation: ^4^.
   ii. n = 1; whole genome sequencing data of DNA extracted from faecal samples. The original study protocol was approved by the Ethical Review Committee, Faculty of Medicine, University of Peradeniya, Sri Lanka [Ref: 2015/EC/58]; citation: this study. Key collaborators: Cinzia Cantacessi, Timothy P Jenkins.
x. Tanzania (n = 5).
   i. n = 5; whole genome sequencing data of DNA extracted from concentrated eggs from faecal samples; ENA study accession: PRJEB44010; sample accession codes: ERR9805806-810; citation: ^2^.
y. Thailand (n = 15).
   i. n = 15; whole genome sequencing data of DNA extracted from individual *S. stercoralis* worms; ENA study accession: PRJNA602131; accession codes: SRR10915458-5472; Citation: ^8^.
z. Uganda (n = 47)
   i. n = 32; whole genome sequencing data of DNA extracted from faecal samples (4 Gb, for 31 samples, 12 Gb for BLANK sample); the original study protocol was approved by the UVRI Research Ethics Committee, as well as the Uganda National Council for Science and Technology and the University of Manchester Research Ethics Committee^9^. Key collaborators: Emma Houlder, Andrew MacDonald, Harriet Mpairwe.
   ii. n = 1; whole genome sequencing data of DNA extracted from a *T. trichiura* worm; ENA study accession: PRJNA304165; sample accession code: SRR2968131; citation: ^3^.
   iii. n = 12; whole genome sequencing data of DNA extracted from *T. trichiura* worms; ENA study accession: PRJEB44010; sample accession codes: ERR9805811-22; citation: ^2^.
   iv. n = 2; whole genome sequencing data of DNA extracted from *S. mansoni* worms; ENA study accession: PRJEB2679; sample accession code: ERR119615; citation: ^1^.
aa. USA (n = 1).
    i. n = 1; whole genome sequencing data from *S. stercoralis*; ENA study accession: [PRJEB2679](https://www.ebi.ac.uk/ena/browser/view/PRJEB2679); sample accession code: ERR066168; citation: ^10^.

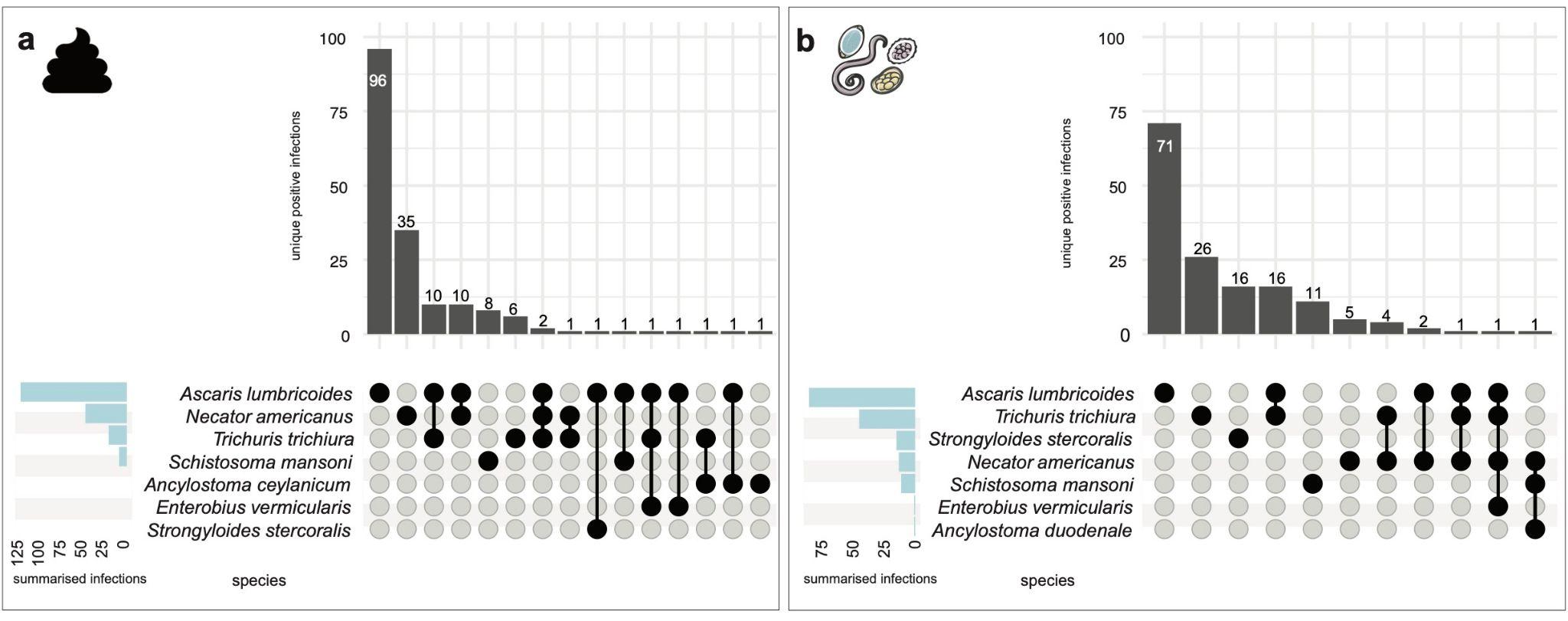

### Supplementary Figure 1: Comparison of helminth-positive samples with single or multiple helminth species infections. The number of single and multiple helminth infections found in a, faecal samples and b, worm/concentrated egg samples. Raw reads were normalised by the total number of reads per sample per genome size to obtain ‘reads mapped per million reads per Mb’; samples were defined as helminth positive if they contained a normalised read count >10. The total number of positive infections is shown (grey vertical bars). The total number of positive infections for each species, summed across all samples, is shown in the blue horizontal bars. The faecal sample icon indicates faecal samples and adult worm/egg figures indicate samples from adult worms and/or concentrated worm eggs.

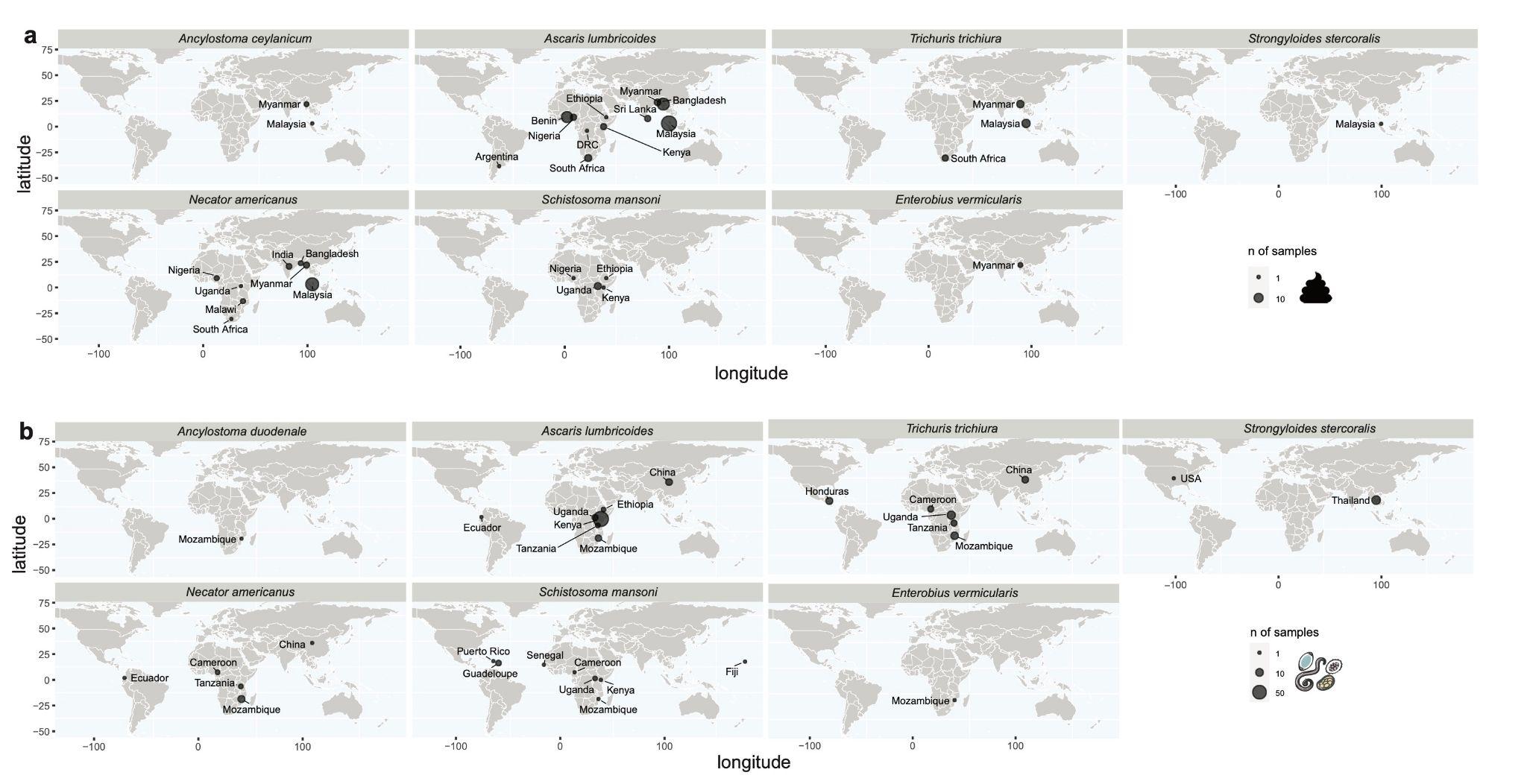

### Supplementary Figure 2: Geographic distribution of helminth-positive samples from faecal and worm isolates.

World maps show the approximate sampling or data origin of **a,** helminth-positive faecal samples (n = 175) and **b,** worm samples (n = 154) detected and analysed in the study. The size of the points on the map is proportional to the number of samples from that location. Raw reads were normalised by the total number of reads per sample per genome size to obtain ‘reads mapped per million reads per Mb’; samples were defined as helminth positive if they contained a normalised read count >10.

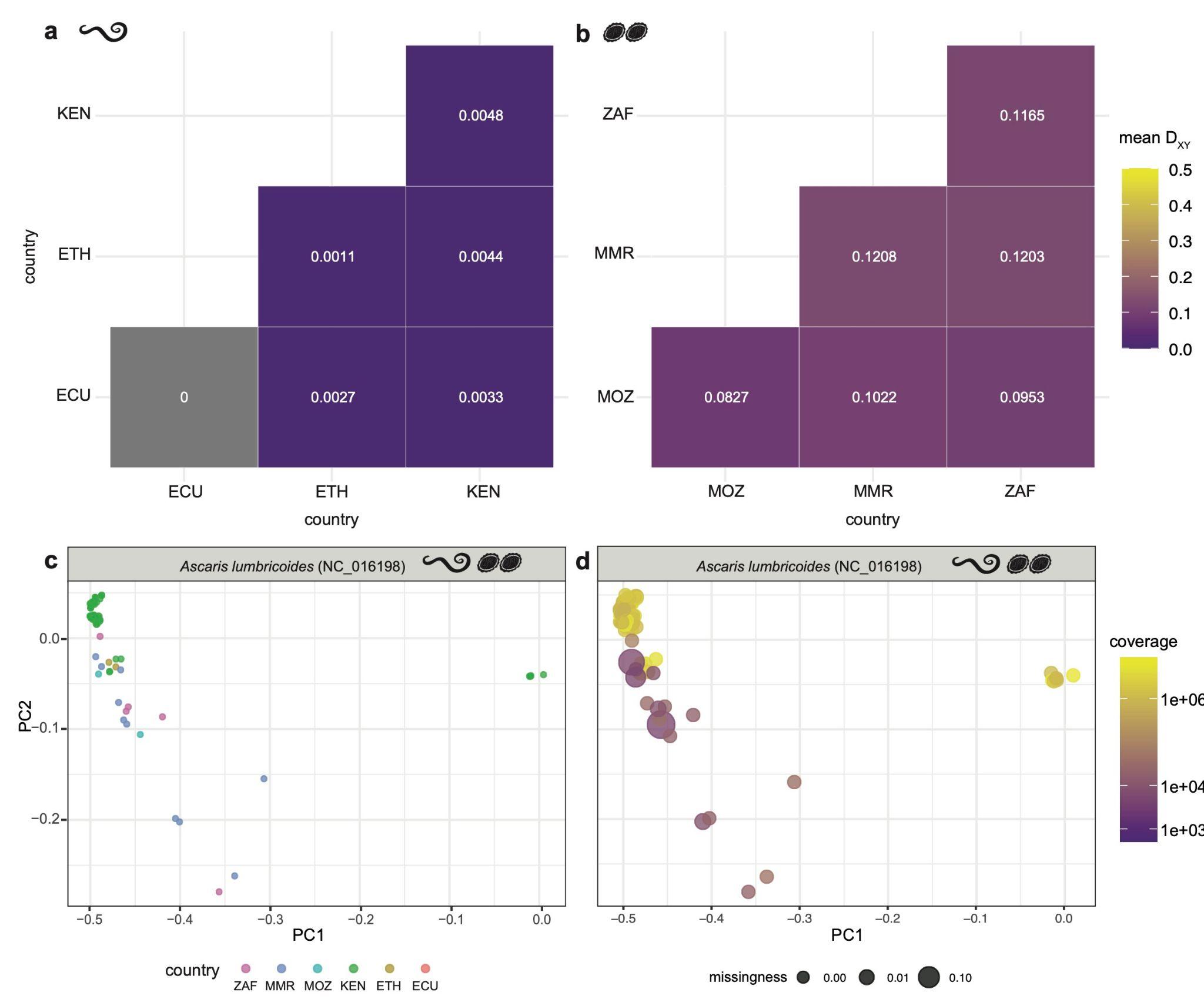

### Supplementary Figure 3: Exploratory analysis of population and genetic structure of samples positive for *Ascaris lumbricoides*.

**a,b,** Heatmaps of pairwise genetic differentiation (D_X_**_Y_**) between *Ascaris lumbricoides* country-specific populations based on pairwise estimates of mitochondrial genome diversity for both worm (a) and pooled (b) samples mapped to the *A. lumbricoides* mitochondrial genome NC_016198 from Korea. The mean D_XY_ per pairwise comparison is shown. **a,** D_XY_ was determined for individual worms using *pixy* on a VCF file containing all filtered variants for all populations. This analysis showed higher variation within Kenya (D_X_**_Y_** = 0.0048) relative to between-country comparisons, such as between Kenya and Ethiopia (D_X_**_Y_** = 0.0044) or between Kenya and Ecuador (D_X_**_Y_** = 0.0033) or Ecuador and Ethiopia (D_X_**_Y_** = 0.0027). Within Ethiopia, the analyses showed that there was little within-country genetic differentiation. **b,** In the pooled samples, D_XY_ was calculated using *Grenedalf* using bam files as input. These analyses showed that there was little genetic differentiation between the countries: Mozambique-South Africa (D_XY_ = 0.0953), South Africa-Myanmar (D_X_**_Y_** = 0.1203), Mozambique-Myanmar (D_X_**_Y_** = 0.1022), Mozambique-Myanmar (D_X_**_Y_** = 0.1022). These comparisons showed very little genetic differentiation between the countries. **c,** Population structure analysis using Bayesian principal component analysis (BPCA) of *A. lumbricoides* (88 samples, 558 SNPs; PC1 = 88.1%, PC2 = 2.1%) identified a few genetically defined clusters, clearly separating most samples from Kenya, whereas samples of mixed origin formed subclusters (South Africa-Myanmar, Myanmar-Mozambique-South Africa, Ethiopia, Kenya, Myanmar). **d,** The BPCA plot for *A. lumbricoides,* as in (c), is coloured by normalised coverage, and points scaled by the degree of missingness are shown. Country codes are as follows: ECU = Ecuador; ETH = Ethiopia; KEN = Kenya; MMR = Myanmar; MOZ = Mozambique; ZAF = South Africa.

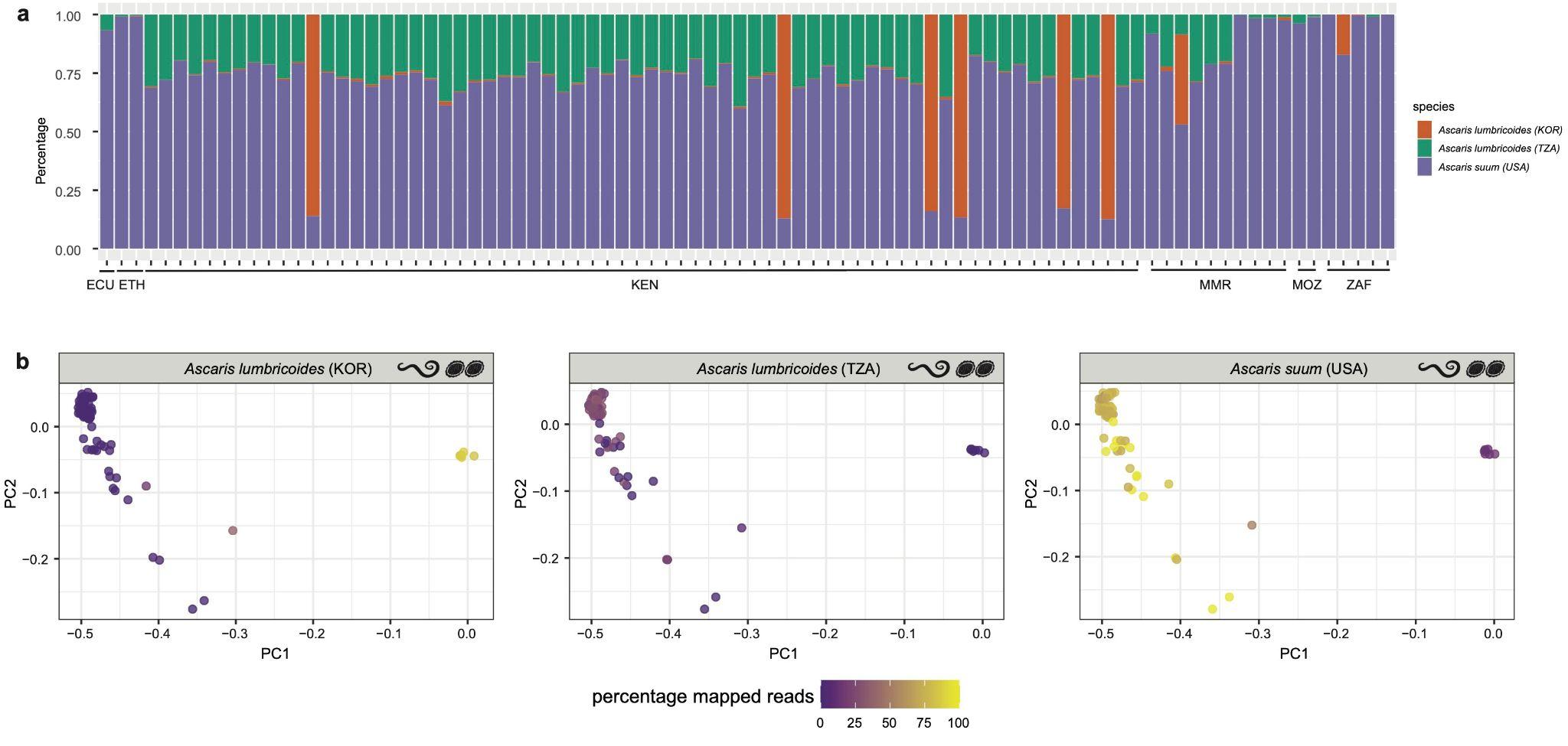

### Supplementary Figure 4. Preferential mapping of samples positive for *Ascaris* spp. to one of three *Ascaris* reference genomes from geographically distinct isolates.

**a,** Samples included in the mitochondrial genome analysis of *Ascaris* spp. were mapped competitively to three different *Ascaris* spp. references, (i) *Ascaris lumbricoides* human isolate (NC_016198, Korea), (ii) *A. lumbricoides* human isolate (KY045802, Tanzania) and (iii) *A. suum* pig isolate (NC_001327, USA). Reads from six samples from Kenya mainly mapped to the human-derived *A. lumbricoides* reference sequence from Korea. In contrast, the remaining samples (n=82) from Ecuador, Ethiopia, Mozambique, Myanmar and South Africa preferentially mapped to *A. suum* (USA isolate). **b,** The PC1 and PC2 components from the BPCA plot of *Ascaris lumbricoides* (NC_016198) (see Supplementary Fig. 3) were re-plotted and colour-coded according to the three different references listed above (*A. lumbricoides* human isolate [NC_016198, Korea], *A. lumbricoides* human isolate [KY045802, Tanzania] and *A. suum* pig isolate [NC_001327, USA]) to reveal any outliers. The outliers (n = 6) that mapped preferentially to the *A. lumbricoide*s isolate from Korea (NC_016198, Korea) were excluded from downstream population analyses. Country codes are as follows: ECU = Ecuador; ETH = Ethiopia; KEN = Kenya; MMR = Myanmar; MOZ = Mozambique; ZAF = South Africa.

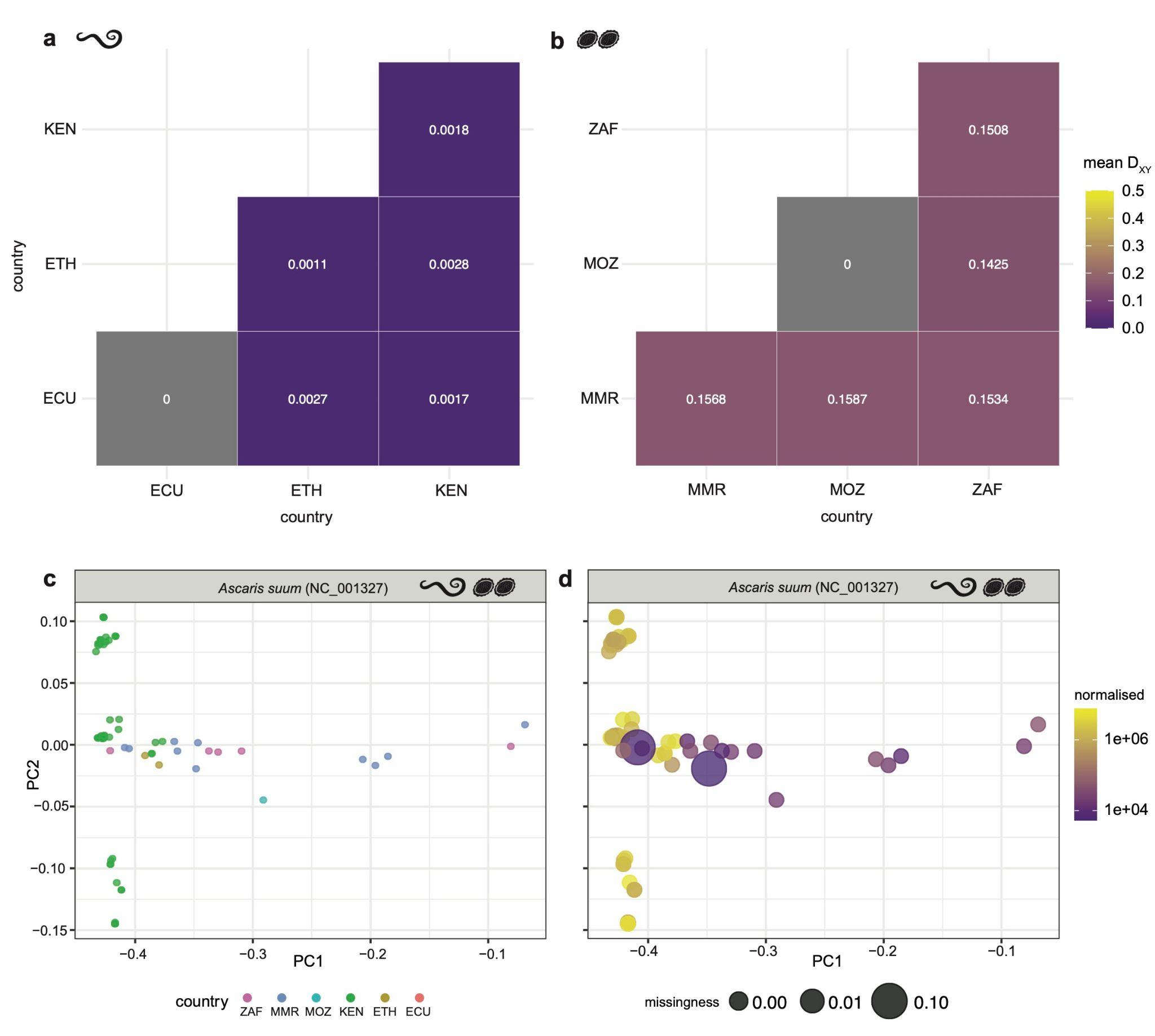

### Supplementary Figure 5. Resolved genetic analyses of *Ascaris* spp. positive samples mapping to *Ascaris suum*.

**a,b,** Heatmaps of pairwise genetic differentiation (D_XY_) between country-specific populations based on pairwise estimates of mitochondrial genome diversity for both worm (a) and pooled (b) samples mapped to the *Ascaris suum* mitochondrial genome NC_001327 from USA. The mean D_XY_ per pairwise comparison is shown. **a**, D_XY_ was determined for individual worms using pixy on a VCF file containing all filtered variants for all populations. The genetic distance between Ethiopia and Ecuador (D_XY_ = 0.0027) and within Ethiopia (D_XY_ = 0.0011) remained unchanged from the original mapping (Supplementary Figure 3). The genetic distance between Ecuador and Kenya decreased (D_XY_ = 0.0017) when samples were mapped to the *A. suum* reference. **b**, In the pooled samples, D_XY_ was calculated using Grenedalf using bam files as input. Heatmap of D_XY_ comparing pooled samples shows that the genetic variation within countries increased relative to the previous analysis (D_XY_ = 0.1508 within South Africa; D_XY_ = 0.1568 within Myanmar), alongside D_XY_ values between countries; D_XY_ = 0.1534 for Myanmar-South Africa (increased); D_XY_ = 0.1425 Mozambique-South Africa (increased); D_XY_ = 0.1587 Mozambique-Myanmar (increased). However, between-country diversity was higher than within-country diversity for Myanmar and South Africa when samples were mapped to A. lumbricoides. For samples from Mozambique, only one sample was retained after filtering for SNP quality and missingness filtering (from *A. suum* mapping), so we could not calculate pairwise within-country diversity. **c,** Population structure analysis using Bayesian principal component analysis (BPCA) of *A. suum* (81 samples, 346 SNPs; (PC1 = 82.4%, PC2 = 3.1%) on samples from Ecuador, Ethiopia, Kenya, Mozambique, Myanmar and South Africa identified high similarity between all Kenyan samples (PC2). The cluster of samples from Ethiopia is closer to the Kenyan cluster (closer geographically), followed by clusters of Myanmar, South Africa, and Mozambique (n=1). The lack of group clustering for all other countries/populations suggests that the primary sources of variation captured by the principal components are not strongly associated with country differences. **d**, The BPCA plot for *A. suum*, as in (c), is coloured by normalised coverage, and points scaled by the degree of missingness are shown. Country codes are as follows: ECU = Ecuador; ETH = Ethiopia; KEN = Kenya; MMR = Myanmar; MOZ = Mozambique; ZAF = South Africa.

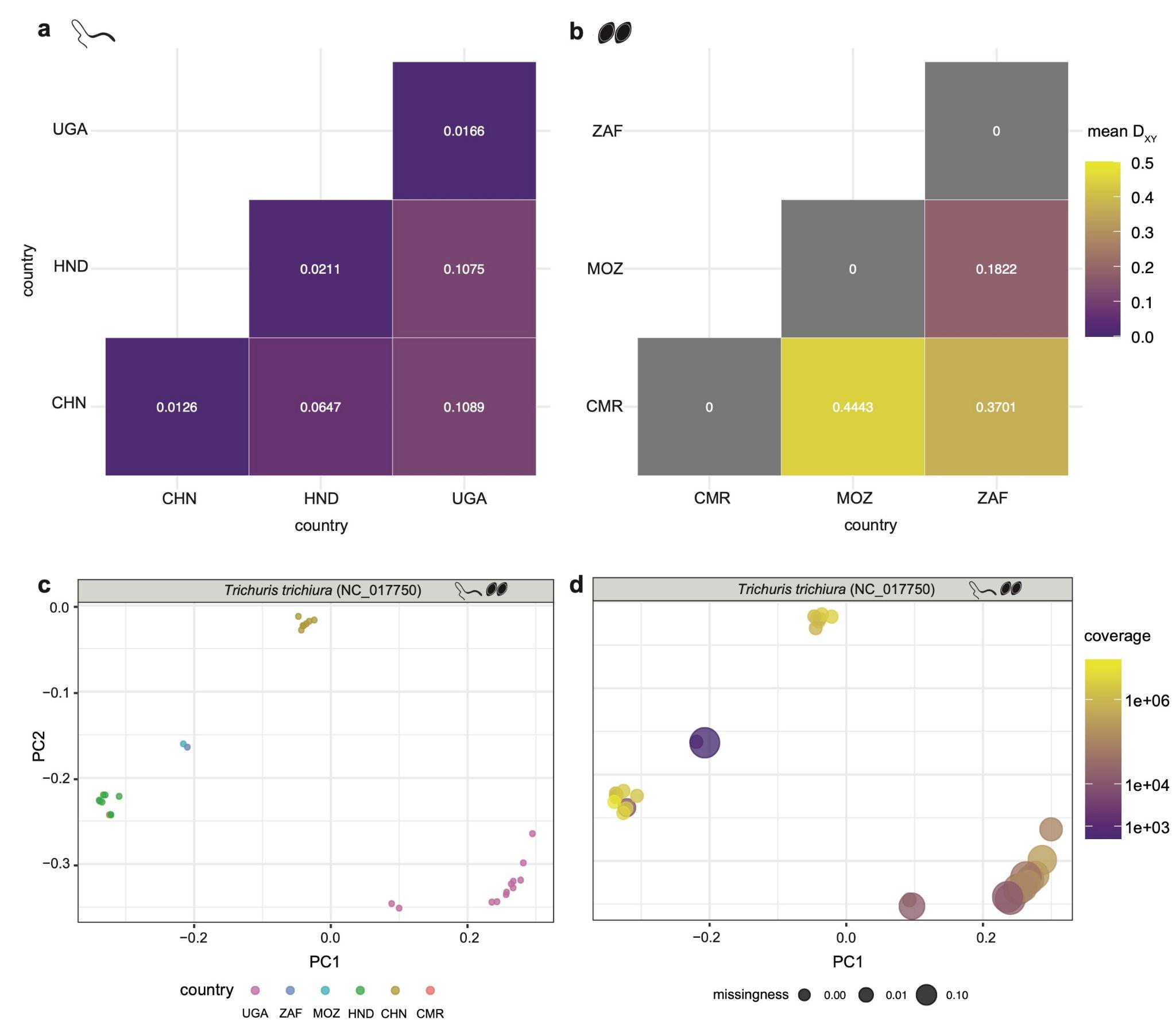

### Supplementary Figure 6. Population structure and genetic differentiation between *Trichuris trichiura* populations.

**a,b,** Heatmaps show pairwise estimates of mitochondrial genome diversity between *Trichuris trichiura* country-specific populations based on pairwise estimates of mitochondrial genome diversity for both worm (a) and pooled (b) samples mapped to the T. trichiura mitochondrial genome from China. The mean D_XY_ per pairwise comparison is shown. **a**, D_XY_ was determined for individual worms using *pixy* on a VCF file containing all filtered variants for all populations. For individual worms within-country variation was lower (Uganda-Uganda: D_XY_ = 0.0166, Honduras-Honduras: D_XY_ = 0.0211, China-China: D_XY_ = 0.0126), compared to between-country variation: Uganda-China more diverse (D_XY_ = 0.1089) than Uganda-Honduras (D_XY_ = 0.1075), and China-Honduras appearing more genetically similar (D_XY_ = 0.0647). **b**, In the pooled samples, D_XY_ was calculated using *Grenedalf* using bam files as input. Heatmap of D_XY_ comparing pooled samples showed that the Cameroon population appears to be more similar to South Africa (D_XY_ = 0.3701) than Mozambique (D_XY_ = 0.4443), but Mozambique seems to be less genetically dissimilar from South Africa (D_XY_ = 0.1822), consistent with geographical distance (Fig. 2a). Overall, the results from genetic distance showed that *Trichuris trichiura* follows anticipated patterns of geographical radiation. **c**, Population structure analysis using Bayesian principal component analysis (BPCA) of *T. trichiura* (30 samples, 1,496 SNPs; PC1 = 40.3%, PC2 = 33.9%) on samples from China, Honduras, Cameroon, Uganda, South Africa and Mozambique. Overall, distinct country-clustering was observed for *T. trichiura* (China, Uganda, Honduras). A mixed cluster of Cameroon (n = 1) and Mozambique (n = 1) with South Africa (n = 1) was also observed. **d**, The BPCA plot for T. trichiura, as in (c), is coloured by normalised coverage, and points scaled by the degree of missingness are shown. County codes are as follows: CHN = China; CMR = Cameroon; HND = Honduras; MOZ = Mozambique; UGA = Uganda; ZAF = South Africa.

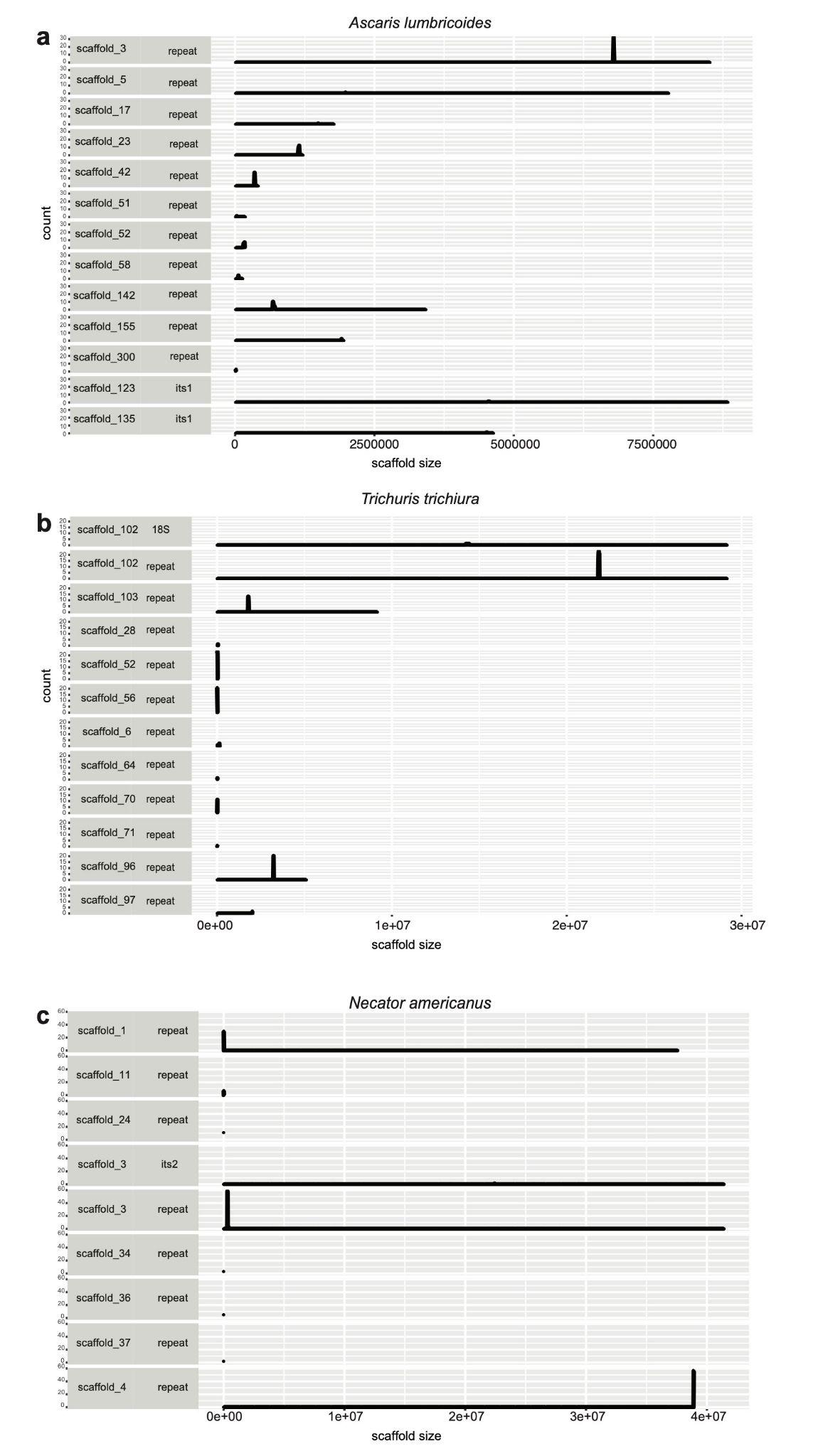

### Supplementary Figure 7. Distribution of diagnostic targets (nuclear repeats, nuclear ribosomal operon genes) in genome assemblies.

Canonical/published repeats were detected using *Nucmer*, allowing for 90% similarity and 90% coverage against the genome assemblies of **a,** *Ascaris lumbricoides* (ALgV5), **b,** *Trichuris trichiura* (trichuris_trichiura_v2), and **c,** *Necator americanus* (Nec_am_Ar_1.0) (Supplementary Table S2). In each plot, the length of the scaffold is shown on the x-axis and counts of repeats per 10 Kb window are shown on the y-axis showing they are tandemly arranged.

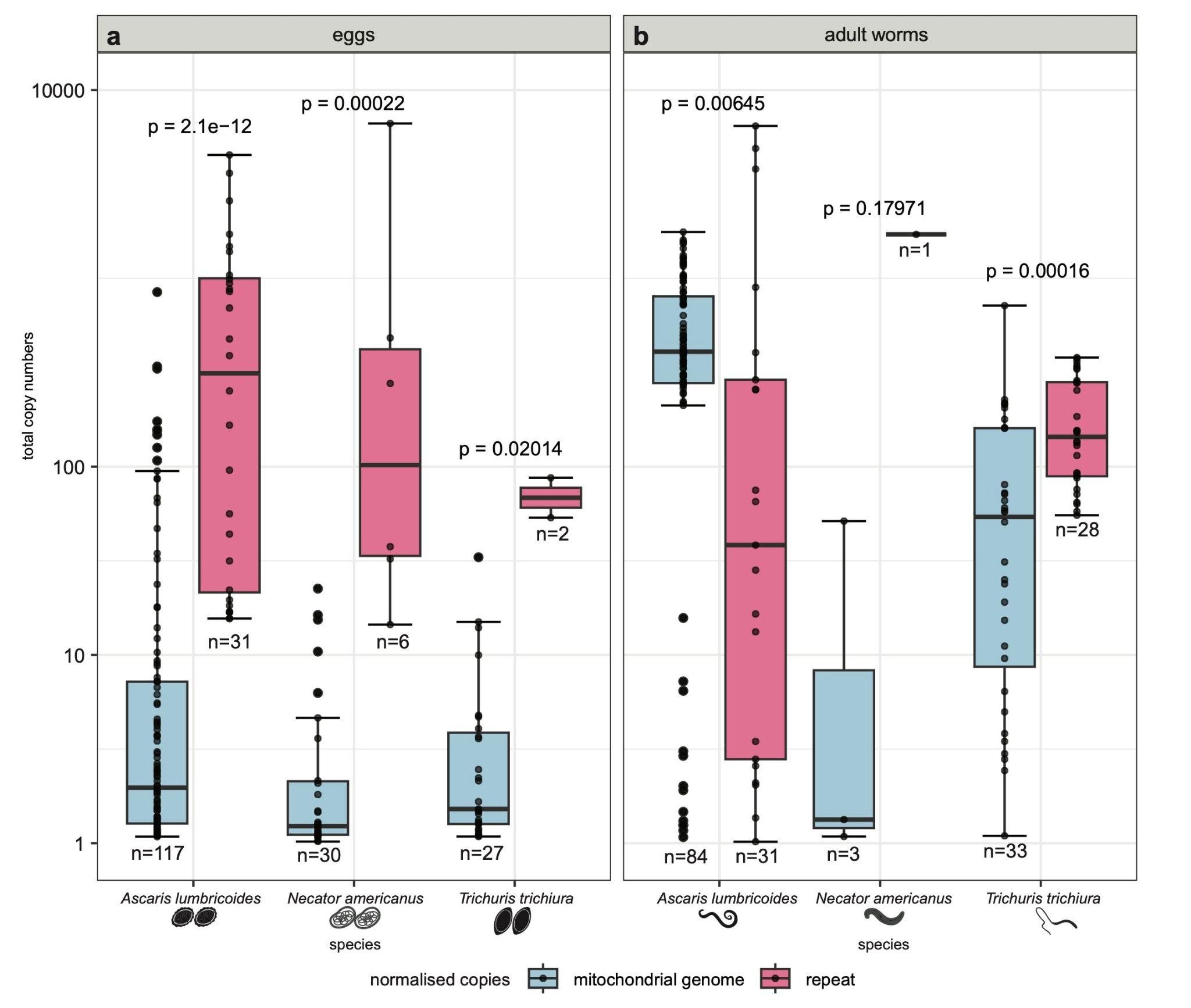

### Supplementary Figure 8. Comparison of relative copy-number of nuclear repeats and mitochondrial genomes based on sequencing coverage.

Boxplots compare the relative copy number of mitochondrial genomes (blue) and repeat units (in pink) per individual per species for **a,** eggs and **b,** adult worms. The relative copy number was calculated per individual by generating a ratio of mitochondrial or repeat coverage divided by the mean coverage of exons from single-copy genes. Each plot shows the sample numbers for each group and the p-value generated from a Kruskal-Wallis test to compare mitochondrial genome copies to nuclear repeat copies per species per parasite developmental stage (with p < 0.05 considered statistically significant). The central box represents the interquartile range and the whiskers represent the data's first and fourth quartile. The median is shown as a line through the centre of the box. The whiskers extend from the edges of the box to the smallest and largest values within 1.5 times the IQR from Q1 and Q3, respectively. Any data points above the upper whisker are considered outliers.

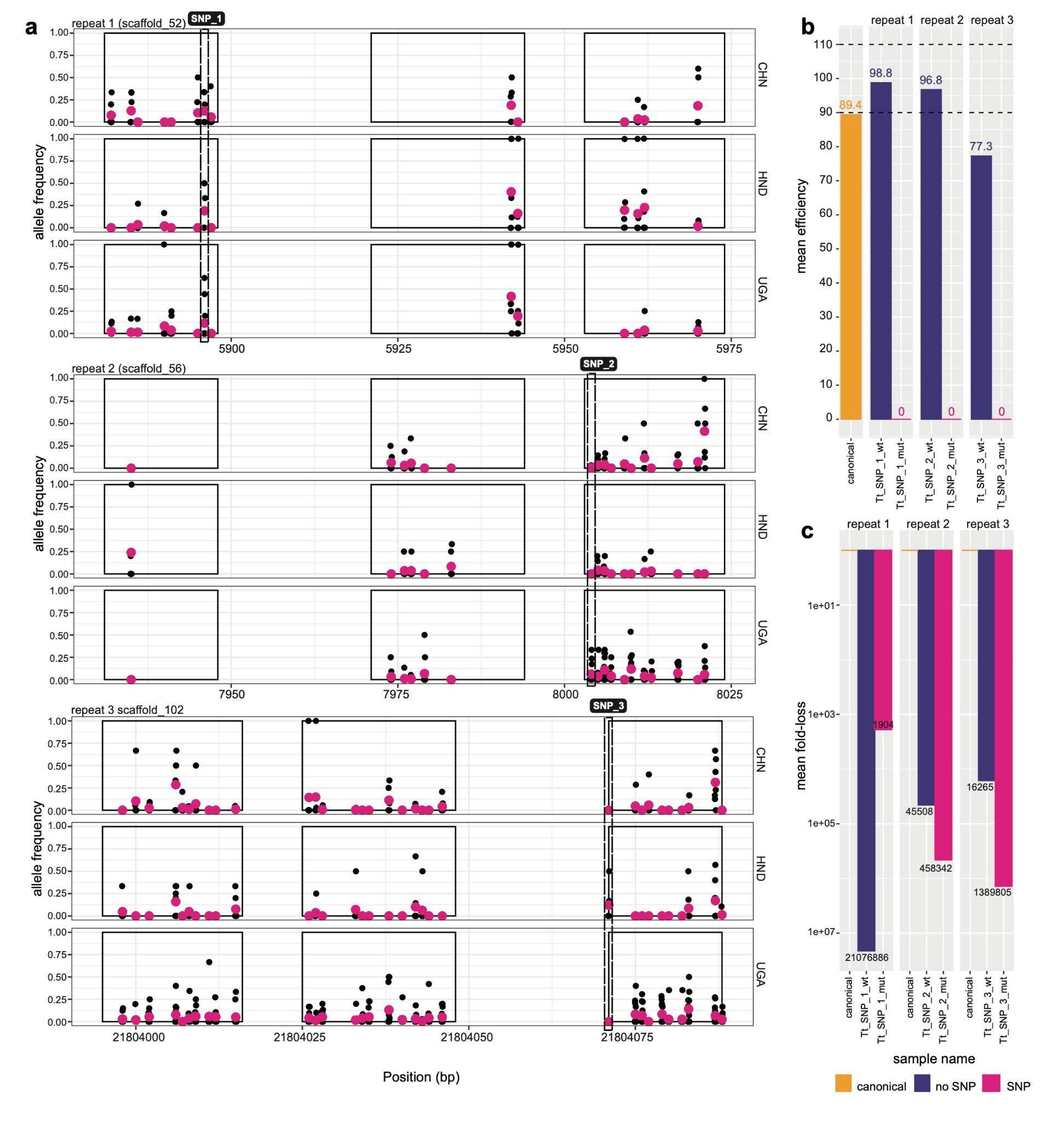

### Supplementary Figure 9. Presence, distribution, and impact of genetic variation within diagnostic qPCR targets of *Trichuris trichiura*.

**a,** Shown are the genomic coordinates of three repeats - repeat 1, 2, and 3 - highlighting primer and probe binding sites (solid rectangles) used in the qPCR diagnostic test, the position of genetic variants (x-axis) - either individual samples (black points) or mean across samples (pink points) - and their frequency within each country (y-axis). Putative qPCR-disruptive variants found at the 3’ end of the primer binding sites are depicted by dashed rectangles. **b,** qPCR efficiencies were determined by generating standard curves of five serial dilutions (100 pg/μl to 10 fg/μl) on each of the repeats in the absence (“wt”) or presence of the SNP (“mut”). The 90-110% dashed lines show the acceptable qPCR efficiency cutoffs. **c,** The mean fold-loss was calculated to assess the effect of the SNP in qPCR quantitation and product loss due to late amplification. Due to the significant effects of the SNPs and other mismatches within the primer binding site of the wildtype samples, the mean fold loss of the wildtype is relative to the canonical repeat. In contrast, the mean fold loss of the mutant is relative to the wildtype repeat within each assay. The mean normalised C_q_ difference was estimated from all serial dilutions. Country codes are as follows: CHN = China; HND = Honduras; UGA = Uganda.
